# Elevation of NAD+ by nicotinamide riboside spares spinal cord tissue from injury and promotes locomotor recovery

**DOI:** 10.1101/2023.01.17.524307

**Authors:** Mariajose Metcalfe, Brian T. David, Brett C. Langley, Caitlin E. Hill

**Author notes:** co-senior authors (Brett C. Langley, Caitlin E. Hill). Reeve-Irvine Research Center, Room 1107, Gillespie Neuroscience Research Facility, 837 Health Sciences Rd., University of California at Irvine, Irvine, CA 92697-4292. Rush University Medical Center, 1725 W. Harrison Street, #855, Chicago, IL 60612. Te Huataki Waiora - School of Health, The University of Waikato, Private Bag 3105, Hamilton 3240, New Zealand, (Brett C. Langley). Neural Stem Cell Institute, One Discovery Drive, Rensselaer, NY, 12144. ^*^**Corresponding author:** Caitlin E. Hill, Principal Investigator, Neural Stem Cell Institute, One Discovery Drive, Rensselaer, NY, 12144. **5. Author contributions:** B.C.L initially conceptualized the project. C.E.H. and B.C.L developed the project and generated the experimental design. C.E.H, M.M, and B.T.D performed the research and analyzed the data; C.E.H., wrote the manuscript. C.E.H, M.M, B.T.D and B.C.L revised and commented on the manuscript.

## Abstract

Spinal cord injury (SCI)-induced tissue damage spreads to neighboring spared cells in the hours, days and weeks following injury leading to exacerbation of tissue damage and functional deficits. Among the biochemical changes is the rapid reduction of cellular nicotinamide adenine dinucleotide (NAD^+^), an essential coenzyme for energy metabolism and an essential cofactor for non-redox NAD^+^-dependent enzymes with critical functions in sensing and repairing damaged tissue. NAD^+^ depletion propagates tissue damage. Augmenting NAD^+^ by exogenous application of NAD^+^, its synthesizing enzymes or its cellular precursors mitigates tissue damage. Among the NAD^+^ precursors, nicotinamide riboside (NR) appears to be particularly well-suited for clinical translation. It safely and effectively augments cellular NAD^+^ synthesis in a variety of species, including rats and humans, and in a variety of preclinical models, elicits tissue protection. Evidence of NR’s efficacy in the context of SCI repair, however, is currently lacking. These studies tested the hypothesis that administration of NR can effectively enhance NAD^+^ in the injured spinal cord and that augmenting spinal cord NAD^+^ protects spinal cord tissue from injury and leads to improvements in locomotor recovery. The results show that intraperitoneal administration of NR (500 mg/kg), administered four days prior to and two weeks following a mid-thoracic contusion-SCI injury, doubles spinal cord NAD^+^ levels in Long-Evans rats. NR administration preserves spinal cord tissue after injury including neurons and axons, as determined by gray and white matter sparing, and enhances motor function, as assessed by the BBB subscore and missteps on the horizontal ladderwalk. Collectively, the findings demonstrate that administration of the NAD^+^ precursor, NR, to elevate NAD^+^ within the injured spinal cord mitigates the tissue damage and functional decline that occurs following SCI.

**HIGHLIGHTS:**

- Nicotinamide Riboside augments spinal cord nicotinamide adenine dinucleotide (NAD^+^).
- Elevating NAD^+^ protects spinal cord tissue from spinal cord injury (SCI).
- Elevating NAD^+^ enhances motor recovery following SCI.

## INTRODUCTION

Damage to the spinal cord triggers a series of secondary events, including pathological changes that perpetuate and spread tissue damage beyond the initial injury (Hall and Springer, 2004; Witiw and Fehlings, 2015). The acute (hours to days) and subacute (days to weeks) post-spinal cord injury (SCI) intervals, during which tissue damage continues and evolves provide an early temporal window of opportunity to limit the spread of tissue damage, enhance tissue sparing, and mitigate functional loss (Ahuja et al., 2017). Currently there are no clinically approved interventions that effectively preserve tissue and limit the extent of functional loss during the early post-injury window (Joaquim et al., 2020; Nagoshi and Fehlings, 2015). As a result, there is an ongoing need to develop and test therapeutic agents that counteract the biochemical mechanisms that exacerbate the tissue damage and lead to SCI-induced dysfunction.

Among the biochemical mechanisms linked to tissue damage is depletion of nicotinamide adenine dinucleotide (NAD^+^) – a key cellular metabolic factor necessary for cell preservation (reviewed in (Covarrubias et al., 2021)). NAD^+^ has several key functions within cells (Covarrubias et al., 2021), including essential roles in basic cellular energy metabolism and as a substrate or co-factor for hundreds of enzymes (Sahar et al., 2011). As part of cellular metabolism, NAD^+^ and its derivatives function as co-enzymes for oxidoreductases and dehydrogenases (i.e., NAD^+^/NADH; NADP^+^/NADPH (nicotinamide adenine dinucleotide phosphate)) (Covarrubias et al., 2021; Srivastava, 2016). It is important for glycolysis, the citric acid cycle, and the mitochondrial electron transport chain, and is needed for the generation of adenosine triphosphate (ATP). NAD^+^ is also a substrate for NAD-consuming enzymes, which include poly-adenosine diphosphate (ADP)-ribose polymerases (PARPs), sirtuins, cyclic-ADP-ribose synthases (e.g., CD38, CD157), and sterile alpha and TIR motif containing 1 (SARM1) (Elhassan et al., 2017; Essuman et al., 2018). These enzymes have important nervous system functions, including roles in cell survival, axon degeneration, senescence, proliferation, apoptosis, DNA repair, cell metabolism and tissue and axon regeneration (Covarrubias et al., 2021). Given the plurality of functions to which NAD^+^ contributes, loss of NAD^+^ has widespread effects. This has led to substantial interest in understanding its roles in the context of nervous system injury and disease.

In healthy tissue, NAD^+^ is depleted and replenished several times a day via several synthetic pathways (Covarrubias et al., 2021; Elhassan et al., 2017; Ijichi et al., 1966). The salvage pathway is the most frequently used to replenish NAD^+^. This pathway uses nicotinamide riboside (NR), vitamin B3, or the degradation product of the NADases (i.e., CD38, CD157, SARM1), and nicotinamide (NAM) as substrates (Covarrubias et al., 2021; Elhassan et al., 2017). Several families of enzymes contribute to NAD^+^ synthesis via the salvage pathway, including nicotinamide mononucleotide adenylyltransferases (NMNATs), which regulate the rate-limiting final step in the pathway to convert nicotinamide mononucleotide (NMN) to NAD^+^, and the nicotinamide riboside kinases (NRKs) and nicotinic acid phosphoribosyltransferases (NAMPTs), which convert NR and NAM, respectively, to NMN. In addition to the salvage pathway, NAD^+^ can be synthesized via the *de novo* pathway using tryptophan or by the Preiss-Handler pathway that uses nicotinic acid (NA) (Covarrubias et al., 2021; Elhassan et al., 2017). Collectively these pathways maintain NAD^+^ levels in cells.

NAD^+^ synthesis attenuation is common in injury and disease, including after SCI where NAD^+^ is rapidly depleted (Hayashi et al., 1984). Genotoxic stress, such as occurs in response to oxidative and nitrosative damage following injury, results in rapid consumption and depletion of NAD^+^ (Alano et al., 2010; Neukomm et al., 2017; Ying et al., 2005). Among the deficits associated with low NAD^+^ levels are cell death (Alano et al., 2010; Ying et al., 2005) and axon degeneration (Figley et al., 2021; Gerdts et al., 2015; Gilley and Coleman, 2010; Sasaki et al., 2006), two key cellular changes associated with spinal cord dysfunction following injury.

In a variety of tissue culture and disease models, NAD^+^ supplementation results in cytoprotection (Alano et al., 2010; Alano et al., 2004; Belenky et al., 2007; Brown et al., 2014; Canto et al., 2012; Gong et al., 2013; Hamity et al., 2017; Harlan et al., 2016; Hou et al., 2018; Khan et al., 2014; Liu et al., 2019; Sasaki et al., 2006; Sasaki et al., 2009; Trammell et al., 2016b; Vaur et al., 2017; Xie et al., 2017a; Xie et al., 2017b; Zhang et al., 2016; Zheng et al., 2019). NAD^+^ can be effectively increased in culture and in the nervous system *in vivo* by exogenous application of NAD^+^, NAD^+^ precursors (e.g., NAM, NMN, NR), or supplementation or manipulation of the enzymes involved in NAD^+^ synthesis (e.g., NMNAT, NAMPT) (Brown et al., 2014; Sasaki et al., 2006; Zhou et al., 2015).

Application of NAD^+^ precursors to increase cellular NAD^+^ levels is particularly effective. Their administration attenuates cell death (Hou et al., 2018), reduces lesion volume (Vaur et al., 2017), counteracts astrocyte toxicity and reactivity (Harlan et al., 2016; Hou et al., 2018), reduces inflammation (Hou et al., 2018; Zhang et al., 2016), modulates oxidative stress (Wei et al., 2017), protects axons and reduces axonal dysfunction (Brown et al., 2014; Gong et al., 2013; Hamity et al., 2017; Hou et al., 2018; Kitaoka et al., 2020; Vaur et al., 2017), stimulates neurogenesis (Hou et al., 2018; Zhou et al., 2020), and reduces cell senescence (Zhang et al., 2016). The ability of NAD^+^ augmentation in other injury/disease models to modify these cellular processes, along with demonstration that exogenous administration of NAD^+^ protects neurons against cell death in ischemic SCI (Xie et al., 2017a; Xie et al., 2017b), led us to examine whether elevating spinal cord NAD^+^ could be an effective treatment for SCI.

Administration of the NAD^+^ precursor, NR, is a particularly promising strategy for elevating NAD^+^ clinically. NR, an alternative form of vitamin B3, is a naturally occurring nucleoside in milk (Trammell et al., 2016c), is endogenously synthesized in cells (Ratajczak et al., 2016), and is commercially available as a nutritional supplement. NR robustly enhances NAD^+^ levels *in vitro* (Chi and Sauve, 2013; Yang et al., 2007; Zhang et al., 2016) and *in vivo* in mice (Canto et al., 2012; Gong et al., 2013; Harlan et al., 2016; Khan et al., 2014; Vaur et al., 2017; Zhang et al., 2016), rats (Hamity et al., 2017), and humans (Trammell et al., 2016a). In preclinical studies, NR is effective at protecting tissue in other models of nervous system injury and disease (Brown et al., 2014; Gong et al., 2013; Hamity et al., 2017; Harlan et al., 2016; Hou et al., 2018; Kitaoka et al., 2020; Vaur et al., 2017; Zhang et al., 2016; Zhou et al., 2020). In humans, NR safely elevates NAD^+^ (Airhart et al., 2017; Conze et al., 2016; Elhassan et al., 2019; Trammell et al., 2016a), and does not appear to have unwanted adverse effects (i.e., painful flushing) that is found with other NAD^+^ precursors (Benyo et al., 2005; Bogan and Brenner, 2008). As NR appears to be safe and effective in other neurological models, and primed for testing and possible clinical translation, in the current study we assessed its effects in a preclinical model of SCI.

This study tests the hypothesis that administration of NR effectively enhances NAD^+^ in the injured spinal cord and that augmenting spinal cord NAD^+^ protects spinal cord tissue from injury and leads to improvements in locomotor recovery. The experimental results reported support the hypothesis. They demonstrate that administration of NR enhances NAD^+^ levels within the injured rat spinal cord, reduces tissue damage, and improves locomotor recovery after a thoracic spinal cord contusion injury in rats. Collectively, the results show that NR is a safe and effective bioavailable agent that can mitigate spinal cord damage.

## MATERIALS AND METHODS

### Experimental Design

Adult (10 – 12-week-old, 230 – 280 g) female Long Evans rats (Charles River Laboratories) were used for the SCI experiments. For preliminary studies assessing the spinal cord biodistribution of NR and its ability to generate NAD^+^, 8-week-old Sprague Dawley rats (Charles River Laboratories) were used. All animal work was performed in accordance with the policies of the Weill Cornell Medicine Institutional Animal Care and Use Committee. For each experiment, the rats were randomly assigned to the treatment groups four days prior to surgery and received either NR or phosphate-buffered saline (PBS) as a control. All *in vivo* experiments were performed and analyzed by investigators blinded to the treatments.

Thirty-nine rats were used in the final assessments of the effects of NR following SCI: NAD^+^ levels (n=10: n=5/group); tissue sparing (n=16: NR, n=7; PBS, n=9), and functional recovery (n=13: NR, n=7; PBS, n=6).

### Drug administration

Purified nicotinamide riboside (NR) was synthesized and provided by Dr. Anthony Sauve (Yang et al., 2007). It was dissolved in sterile PBS and administered intraperitoneally (i.p.) to the rats. In the preliminary studies used to initially determine NR dose, naive rats received one of four different doses of NR (0, 100, 250 and 500 mg/kg/day) for 4 days before tissue was collected, processed, and NAD^+^ quantified. For SCI studies, NR (500 mg/kg/day) or PBS vehicle control was administered to rats beginning 4 days prior to injury. Administration concluded 14 days post-injury (dpi), for a total administration time of 18 days.

### Spinal cord injury

Rats (n=64) received spinal cord injuries along with NR (500 mg/kg/day) or PBS treatment. A complete description of the SCI procedure can be found in (Dai and Hill, 2018). Briefly, rats were anesthetized with isoflurane (3.5% in oxygen) and a 200 kdyn contusion was induced using the Infinite Horizon Impactor (Precision Systems & Instrumentation) following a laminectomy at the 9^th^ thoracic vertebral level (T9). After injury, the muscles were sutured closed in anatomical layers and the animals were given gentamycin (5 mg/kg), buprenorphine (0.05 mg/kg), and lactated Ringer’s solution (10 ml) subcutaneously. Buprenorphine and lactated Ringer’s solution were given twice a day for the first 2 dpi to prevent pain and dehydration, respectively. Gentamycin was given daily for the first 7 dpi to prevent infection. The rats’ bladders were manually expressed twice daily until reflexive urination returned. Rats were provided with food and water *ad libitum*. The survival time of the rats were 14 dpi, 28 dpi, or 9 weeks post-injury (wpi).

### Study exclusion criteria

Rats with a maximum injury force that differed +/− 20% from 200 kdyn, forces that were not continuous upon inspection of the force and displacement curves, BBB scores that differed > 3 points between right and left hindlimbs at either 1 or 7 dpi, or with injuries at the incorrect level (confirmed at time of sacrifice) were excluded. For assessment of NAD^+^ levels following SCI and tissue sparing, 26 rats met the inclusion criteria. From these 26 rats, 5 rats from each group were randomly selected at 14 dpi for sacrifice to examine NAD^+^ levels in the tissues; tissue from the remaining rats was collected at 28 dpi. For assessment of functional recovery, 13 rats met the inclusion criteria and were analyzed for functional recovery.

### Behavioral testing

#### Basso-Beattie-Bresnahan (BBB) locomotor rating scale

For BBB testing, all rats were pre-handled and acclimatized to the testing environment for 10 days before collecting baseline open-field locomotor ability using the 21-point BBB score (Basso et al., 1995) and the 13-point BBB subscore (Basso, 2004) to ensure that all animals had preinjury scores of a 21 and 13, respectively. After SCI, BBB testing was performed on days 1 and 7 for all rats to confirm that the injury was centered. For the 9-week behavioral study, BBB data was collected on days 1, 7, 14, 21, 35, 42, 49, 56, 63, and 70. All BBB testing was performed by two independent, trained observers who were blinded to the treatment groups. BBB scores and BBB subscores are reported as mean ± SEM. For statistical analysis of the BBB and BBB subscore, a two-way mixed model analysis of variance (ANOVA) was used for analysis. For analysis of the proportion of rats with specific subscore components a Chi-squared or Fisher’s exact test was used.

#### Ladderwalk

For the 9-week behavioral study, all rats were pre-trained to cross a horizontal ladder (or the CatWalk on alternate days) for 10 days. Videos of the rats crossing the ladder were acquired prior to drug administration (pre-injury) and every other week starting 2 weeks post injury (wpi). The steps from three passes across the horizontal ladder were analyzed for each rat at each time point. Data was scored using the scale by Metz and Whishaw (Metz and Whishaw, 2002) using the analysis of scorable steps. For ladderwalk statistical analysis, a multivariate repeated measures ANOVA was used.

#### CatWalk

For the 9-week behavioral study, all rats were pre-trained to cross the CatWalk (or horizontal ladder on alternate days) for 10 days. CatWalk data was acquired prior to drug administration (pre-injury) and every other week starting 2 wpi. A total of five passes were collected and assessed at each time point. For statistical analysis, a multivariate repeated measures ANOVA was used.

### Tissue processing

#### NAD^+^ assessment

Fresh frozen tissue was collected for assessment of NAD^+^ levels 4 h after the last i.p. administration. Rats (n=10) were terminally anesthetized with Euthasol and the spinal cord, brain, heart, liver, skeletal muscle and blood were rapidly dissected/collected, flash-frozen on powdered dry ice, and stored at −80 °C. To isolate NAD^+^, the tissue was pulverized in liquid nitrogen, the powder was weighed and 1 M perchloric acid was added and the solution was sonicated. The insoluble fraction was then pelleted and the supernatant neutralized. NAD^+^ levels were determined by a 96-well plate assay utilizing lactate dehydrogenase and diaphorase, as previously described (Yang et al., 2007).

#### Histology

Fixed spinal cord tissue was collected for histological assessment of tissue sparing. Rats were terminally anesthetized with Euthasol and transcardially perfused with heparin and saline, followed with 4% paraformaldehyde in 0.1 M Sorenson’s buffer pH 7.2. Spinal cords were removed, post-fixed overnight, and then cryoprotected in 30% sucrose in 0.1 M PBS. Serial cryostat cross sections (20 μm thick) spanning the injury were cut and thaw-mounted onto electrostatically charged glass slides. To visualize spared tissue (gray matter, white matter and total tissue sparing), slides were stained with BrainStain Imaging Kit (ThermoFisher Scientific), which included NeuroTrace 530/615 red fluorescent Nissl (1:30), FluoroMyelin green dye (1:30), and DAPI (4’, 6-diamidino-2-phenylindole dihydrochloride; 1:30) and cover-slipped with Vectashield mounting medium (Vector Laboratories) prior to examination.

### Microscopy and histological quantification of tissue sparing

Tissue sparing was visualized and assessed 28 days post-SCI (NR: n=9; PBS control: n=7). Tissue sections were examined and imaged using a Zeiss AxioImager M2 with a Hamamatsu C11440 camera. The area of spared gray matter, white matter and total tissue was quantified in every 20^th^ tissue section using the Cavalieri function (100 x 100 μm grid) in Stereo Investigator (MBF Bioscience, version 11.06). Spared gray matter was defined as tissue within intact or damaged dorsal or ventral horns, or central gray matter, with a neuronal morphology (cell body and location) and labeled with red fluorescent Nissl staining. Spared white matter was defined as tissue that had green FluoroMyelin staining. Spared tissue that was neither gray nor white matter (i.e., trabecula) was not included in either estimation, but was included in total tissue sparing. All analyses were conducted by an investigator blinded to the treatment groups. Representative images of tissue sections for figures were acquired using Stereo Investigator. The same settings were used to acquire all images. Two-way mixed model analysis of variance (ANOVA) was used to compare the results of tissue sparing. When appropriate, ANOVAs were followed by Tukey’s post-hoc tests.

### Statistical analysis

All quantitative assessments were performed blinded with respect to the treatment group. IBM SPSS Statistics 22.0 was used to determine the statistical differences between experimental conditions. A p value ≤ 0.05 was considered to be statistically significant. Data are reported as mean ± standard error of the mean (SEM) throughout.

## RESULTS

### The rodent spinal cord expresses the enzymes necessary to synthesize NAD^+^ from NR

The synthesis of NAD^+^ from NR depends on the availability of the NRK1 and NMNATs to convert NR to NMN and NMN to NAD^+^, respectively (Elhassan et al., 2017) (**Fig. 1A**). From development until post-natal day 21, the rat spinal cord expresses the enzymes necessary to generate NAD^+^ (Sasaki et al., 2006). Examination of the Allen mouse spinal cord atlas for expression of NRKs (*Nmrk1* and *2*) and the NMNATs (*Nmnat1, 2, 3*) reveal that expression by all three *Nmnats* are detectable by in situ hybridization (data for *Nmrk1* and *2* are not available). *Nmnat2* shows the greatest expression (**Fig. 1B**, Allen Institute for Brain Science. Mouse Spinal Cord Atlas, mousespinal.brain-map.org). Examination of available single cell expression data on the mouse spinal cord (Milich et al., 2021), showed that *Nmrk1*, and *Nmnat1-3*, but not *Nmrk2* are ubiquitously expressed in various cell types in both the intact (**Fig. 1B**) and acutely and sub-acutely injured rodent spinal cord (**Fig. 1C**). *Nmrk1* and *Nmnat2* had the greatest expression within the spinal cord, both were expressed in neurons, astrocytes, oligodendrocytes progenitor cells and ependymal cells. *Nmrk1* was predominantly expressed in oligodendrocytes, microglia and macrophages, whereas *Nmnat2* showed more limited expression in these cell types (**Fig. 1C**). The examination of the enzymes needed for NAD^+^ synthesis demonstrates that the rodent spinal cord contains the necessary synthetic enzymes to convert NR to NAD^+^.

**Figure 1:**
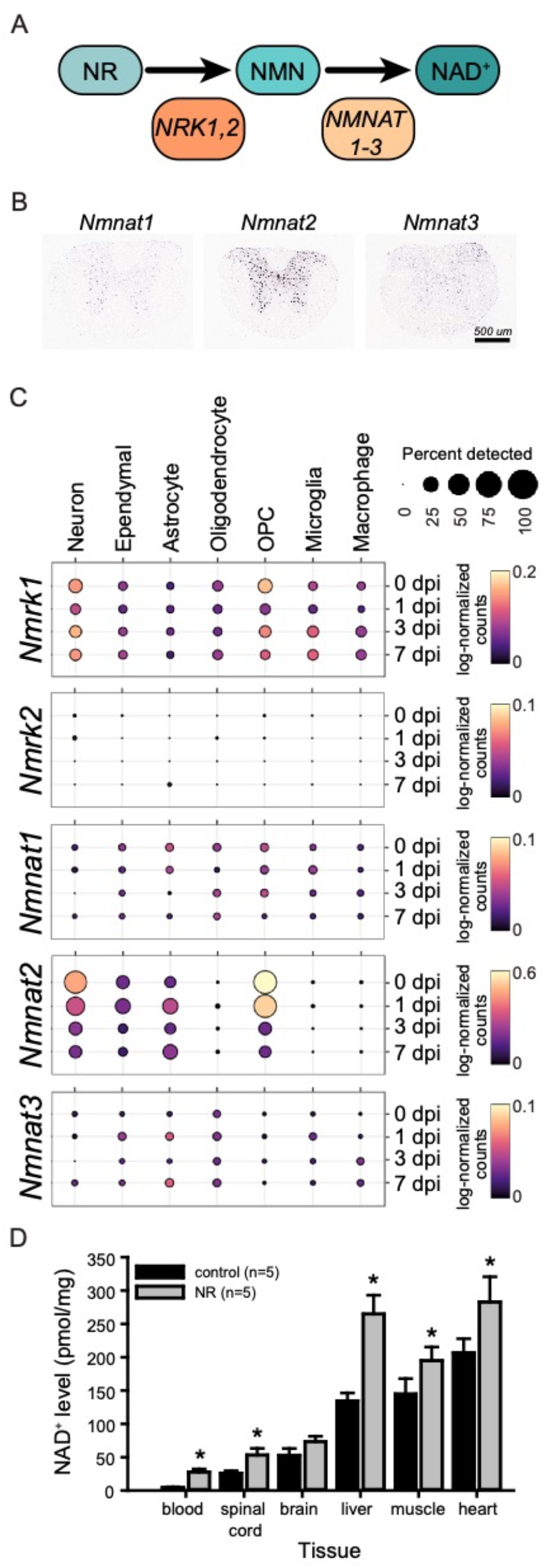
Spinal cord expression of enzymes involved in NAD^+^ synthesis from NR (A-C) and the tissue levels of NAD^+^ following NR administration of SCI rats (D). Schematic showing the enzymes and intermediates needed to convert NR to NAD^+^ **(A)**, the enzyme families involved are the NRKs (gene name *Nmrk*) and the NMNATs. Expression of *Nmnat1–3* in the mouse spinal cord **(B)**. Cell type expression of *Nmrk* and *Nmnat* family members in primary spinal cord cell types within the uninjured and injured mouse spinal cord **(C)**. NAD^+^ levels in tissues isolated from SCI rats following NR administration **(D)**. Administration of 500 mg/kg of NR for 18 days, starting 4 days prior to injury, results in a significant increase in NAD^+^ levels relative to PBS(control) treatment, in a variety of tissues 14 days post-SCI, including the injured spinal cord [F(8,1) = 30.386, one-tailed p=0.0005]. The level of NAD^+^ differed significantly between the different tissues [F(21.59, 2.70) = 67.782, p < 0.0001]. Statistics: Multivariate ANOVA. * denotes p-value of 0.05 or less. Scale = 500 μm.

### Spinal cord NAD^+^ increases with NR administration

To establish that NR was able to cross the blood brain/spinal cord barrier and determine NR dosing, naive rats were treated with different concentrations of NR and the levels of NAD^+^ in the spinal cord were measured in a preliminary study. Compared to saline treated controls, four hours after the terminal injection of NR, spinal cord levels of NAD^+^ were increased by ~ 18%, 35% and 47% following administration of 100, 250, or 500 mg/kg of NR, respectively. NR (250 mg/kg) commencing either 4 days prior to injury or 2 hours post-injury, was initially tested in rats (n =60 rats, n=15 rats/group), however, this dose failed to result in protection of spinal cord tissue or recovery of sensory or motor function (not shown). This led us to test a higher dose of NR. The effects of administering 500 mg/kg NR are presented.

### NR treatment enhances NAD^+^ in multiple tissues, including the injured spinal cord

Following injury, the requirement for NAD^+^-dependent reactions increases and the expression of key NAD^+^ synthetic enzymes changes (**Fig. 1C**)(Sasaki et al., 2006). To test whether 500 mg/kg of NR resulted in an increase in NAD^+^ levels 14 days after spinal cord injury, NR or PBS control was administered for 18 days starting 4 days prior to injury and the level of NAD^+^ was quantified at the SCI site. For comparison, NAD^+^ levels were also quantified in the blood, brain, liver, skeletal muscle and heart. NAD^+^ was significantly increased following NR treatment in all tissues except the brain (p≤0.0005) (**Fig. 1D)**. The overall level of NAD^+^ within the injured spinal cord was substantially lower than the levels detected in the liver, muscle, or heart (**Fig. 1D**). However, administration of NR increased NAD^+^ levels in the spinal cord by 2.0 +/− 0.3-fold, a similar magnitude as was found in the liver (2.0 ± 0.2-fold). The results of the NAD^+^ quantification post-SCI confirmed that 500 mg/kg NR effectively increases NAD^+^ levels within the injured rat spinal cord.

### NR enhancement of NAD^+^ increases tissue preservation following SCI

In other experimental models, increasing, or preserving tissue NAD^+^ levels in tissue are able to counteract excitotoxic damage (Vaur et al., 2017) and oxidative stress (Xie et al., 2017a), reduce cell death (Hou et al., 2018; Xie et al., 2017a; Xie et al., 2017b), prevent axon degeneration (Brown et al., 2014; Gilley and Coleman, 2010; Perry et al., 1991; Perry et al., 1990; Sasaki et al., 2006), and attenuate lesion expansion (Vaur et al., 2017). To assess whether supplementing NAD^+^ is sufficient to preserve spinal cord tissue during the acute/subacute interval after SCI when extensive cell death and tissue degeneration occurs, NR (500 mg/kg) was administered and total tissue sparing was assessed 28 days after injury. **Figure 2A** shows the appearance of NR and PBS (control) treated spinal cords. NR treatment significantly increased spared tissue at the lesion epicenter and throughout a 4-mm segment of spinal cord surrounding the injury epicenter (**Fig. 2B**). Following NR treatment, 24.4% ± 1.2% of tissue was spared at the epicenter compared to 17.1% ± 1.5% in the control. Thus, at 500 mg/kg NR was able to enhance the amount of spared tissue following SCI.

**Figure 2:**
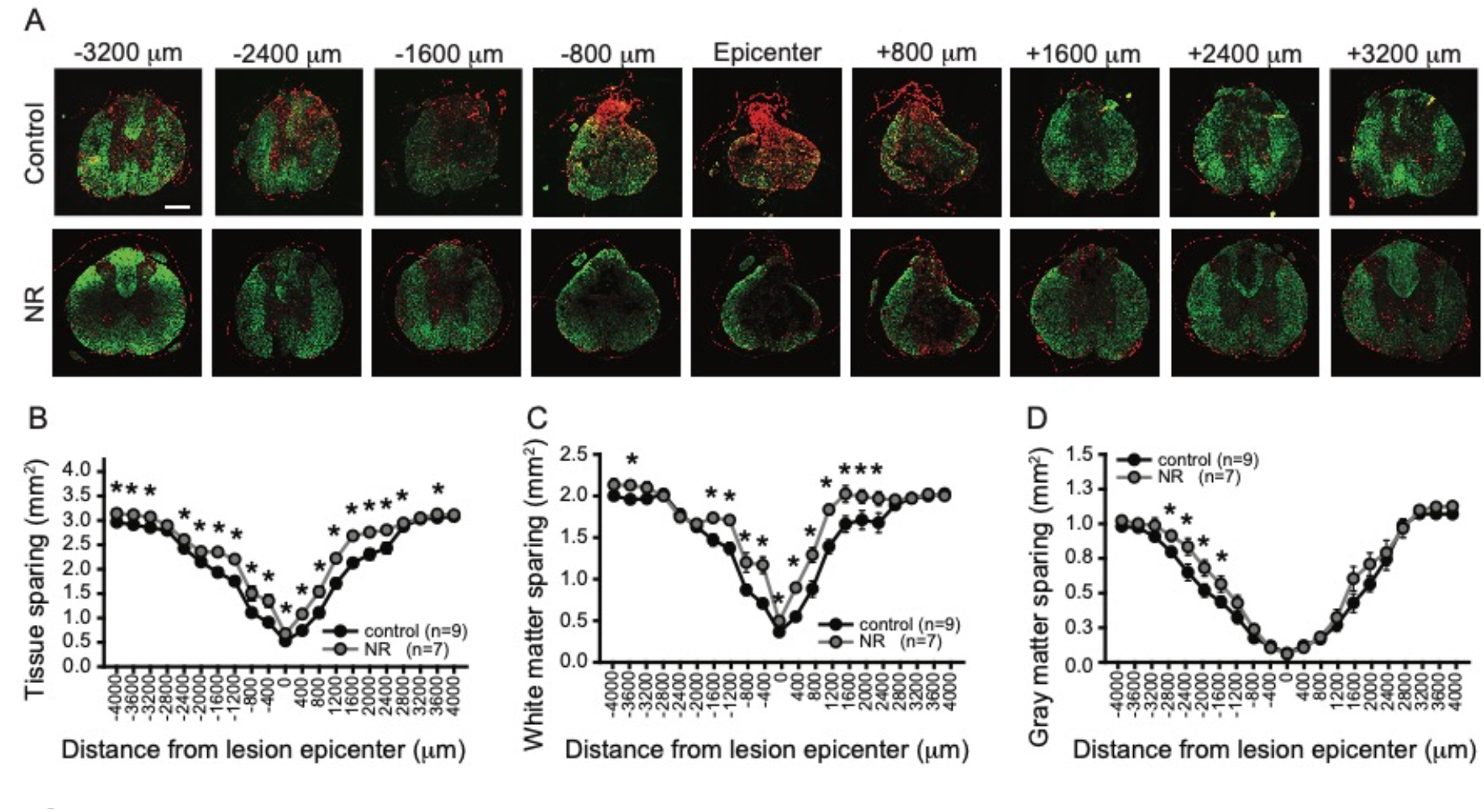
Administration of NR increases tissue sparing 28 days post-SCI. Appearance of spinal cord tissue sections 28 days post-SCI stained with NeuroTrace fluorescent Nissl (red) and FluoroMyelin (green) from −3200 μm rostral to the lesion epicenter to +3200 μm caudal **(A)**. Results of stereological quantification for **B)** total tissue sparing. **(C)** white matter sparing. **(D)** gray matter sparing. Compared to PBS (control) treatment, NR treatment increased total tissue sparing (**B**: F(14,1) = 28.41, one-tailed p < 0.0001), white matter sparing (**C;** F(14,1) = 11.256, one-tailed p = 0.0025) and gray matter sparing (**D**: F(14,1) = 3.40, one-tailed p = 0.043). Statistics: Multivariate ANOVA. * denotes p-value of 0.05 or less. Scale = 500 μm.

### NR enhancement of NAD^+^ attenuates loss of spinal cord white matter following injury

Maintenance of NAD^+^ in axons effectively prevents their degeneration (Brown et al., 2014; Di Stefano et al., 2017; Gerdts et al., 2015; Sasaki et al., 2006; Sasaki et al., 2009; Vaur et al., 2017). To establish whether the tissue sparing observed following NR administration resulted from an increase in axonal sparing, the amount of white matter preserved following injury was examined. NR treatment resulted in more white matter throughout the injury site (**Fig. 2C**). Increased white matter was detected from 1600 μm rostral to 2400 μm caudal to the injury epicenter. At the epicenter, 45% more white matter was present with NR treatment (white matter sparing, NR: 22.9% ± 1.3%; control: 15.8% ± 2.0%). This result indicates that NR elicits tissue protection in part by enhancing the preservation of axons in the injured spinal cord.

### NR enhancement of NAD^+^ preserves neurons at the rostral margin of the injury

In addition to preserving axons, NAD^+^ also enhances cell survival (Hou et al., 2018; Xie et al., 2017a; Xie et al., 2017b). To test whether NR administration protected neurons adjacent to the injury, the area of neurons within the gray matter was quantified. Damage to the spinal cord at thoracic level 9 abolishes virtually all the neurons within the gray matter at the injury epicenter and results in the partial loss of gray matter neurons up to 3200 μm in either direction (**Fig. 2D**). NR treatment increased gray matter area in tissue sections from 1600-2800 μm rostral to the lesion (**Fig. 2D**) but had no effect on the percentage of gray matter sparing at the epicenter (NR: 2.2% ± 1.2%; control: 2.9% ± 0.1%). Thus, in addition to preserving axons, NR treatment augmented neuronal survival within the rostral penumbra of the injury.

### NR enhances functional recovery following SCI

Damage to axon tracts and disruption of the spinal circuitry is a primary contributor to the loss of function following SCI (Taccola et al., 2018). Locomotor recovery correlates with the extent of tissue sparing (Basso et al., 1996) and sparing of as few as 5-10% of axons can result in effective, albeit defective, locomotion (Blight, 1983). To determine whether NR-induced tissue sparing, particularly white matter sparing, was sufficient to promote locomotor recovery, we tested whether administration of NR to rats with a thoracic contusion SCI affected recovery of gross locomotor and fine hindlimb function over the course of 9 wpi when compared to PBS control treatment. In this injury model, disruption of white matter is the primary cause of locomotor deficits (Magnuson et al., 1999).

As expected based on the extent and location of tissue sparing at the epicenter following SCI, both control and NR treated rats showed similar gross locomotor recovery in the open field as assessed by the Basso-Beattie-Bresnahan (BBB) test (Basso et al., 1995), a 21 point scale in which progression of recovery after SCI is assessed (control: 11.7 ± 0.3; NR: 11.9 ± 0.3; **Fig. 3A**). Following injury, rats regained the ability to consistently step, but were not able to regain coordination between the forelimbs (FL) and hindlimbs (HL). The lack of reestablishment of coordination was confirmed by CatWalk gait analysis in which no differences between the groups were detected by the regularity index measurement (not shown). FL-HL coordination is dependent on a diffuse set of axonal pathways located within the ventral and lateral funiculi in the rat spinal cord (Loy et al., 2002a; Loy et al., 2002b). Among the tracts located in these funiculi, propriospinal axons are particularly important for forelimb-hindlimb coordination (Juvin et al., 2005). Few propriospinal axons remain following thoracic contusive SCI (Basso et al., 2002; Conta and Stelzner, 2004) due to their location within the ventrolateral white matter (Reed et al., 2006; Reed et al., 2009), which makes them particularly susceptible to contusive SCI. Even with the increased white matter sparing following NR treatment, the limited sparing of propriospinal axons at the lesion epicenter, likely prevented the reestablishment of forelimb-hindlimb coordination.

**Figure 3:**
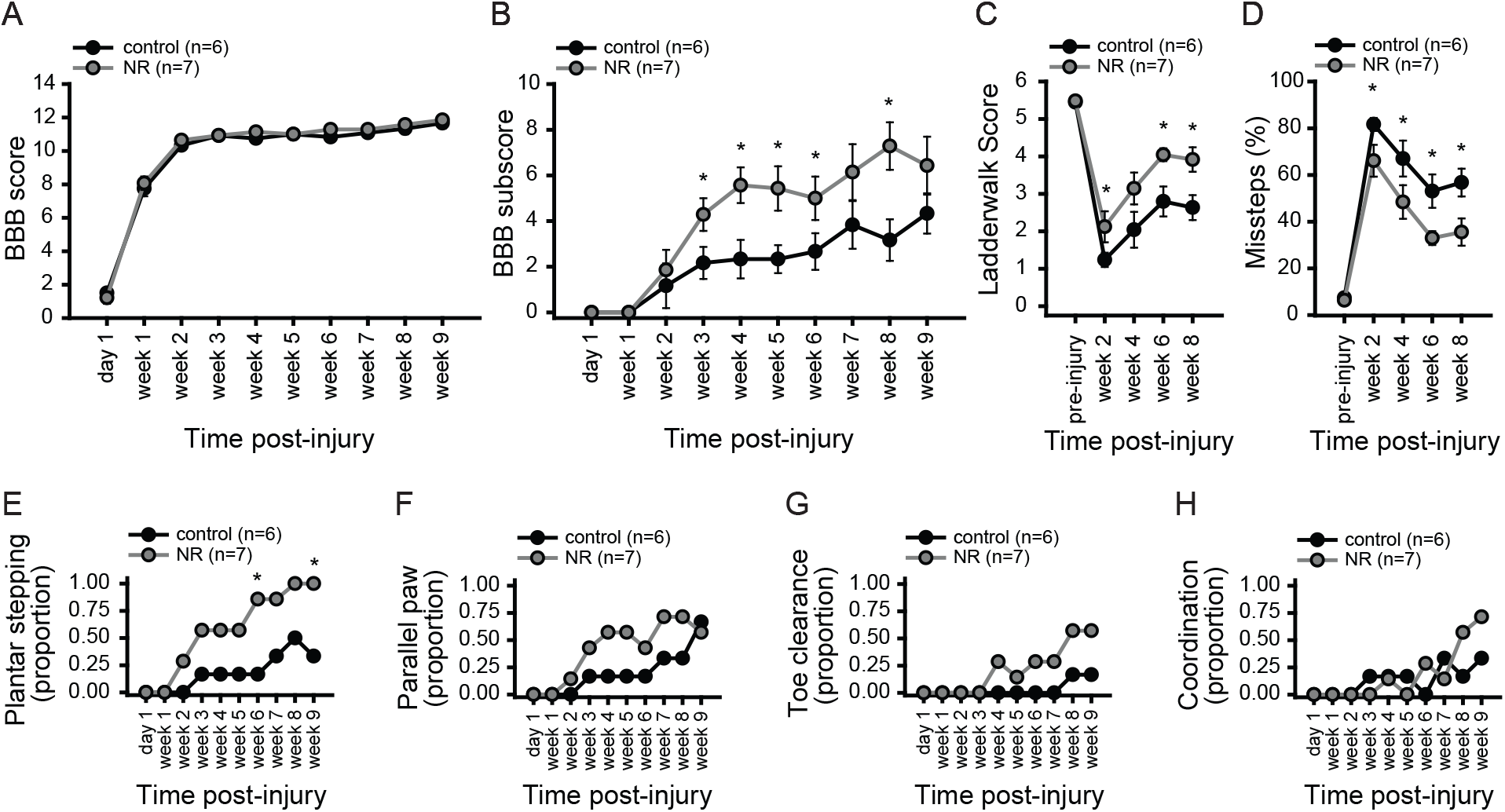
Post-spinal cord injury locomotor function recovers better following NR treatment. Following SCI, both NR and PBS (control) treatment groups had similar gross locomotor recovery, as assessed in the open field with the BBB score **(A)**. Compared to PBS-control treatment, NR treatment resulted in a significant improvement in locomotor recovery as determined by the BBB subscore (**B**: F(11, 1) = 4.29, one-tailed p = 0.032) and the ladderwalk score (**C**: F(11, 1) = 7.35, p = 0.02, one-tailed p = 0.01). NR treated rats made fewer errors on the ladderwalk (**D**: F(11, 1) = 7.86, one-tailed: p < 0.01). Assessment of the proportion of rats demonstrating specific criteria that make up the BBB subscore revealed a greater proportion of rats with NR treatment were able to plantar step (**E**: p = 0.021). Differences between groups on other components that make up the subscore did not differ between NR-treated and control treated rats [proportion of rats with: one or more parallel paws on placement **(F)**; any toe clearance (**G)**; any coordination **(H)**]. Statistics: A-D: Multivariate ANOVA; E-H: Chi squared, Fisher’s exact test. * denotes p-value of 0.05 or less.

The requirement of FL-HL coordination creates a ceiling effect when using the BBB that can mask meaningful functional improvements in hindlimb control. To assess whether NR treatment promoted recovery on aspects of hindlimb function assessed but not scored using the BBB until after coordination is achieved (i.e., consistency of stepping, paw position, toe clearance, trunk stability and tail position) the BBB sub-score was used (Basso et al., 2002). NR treatment resulted in a significant improvement in locomotor recovery using the BBB sub-score, beginning 3 weeks following injury (**Fig 3B**). To determine the specific aspects of recovery that were improved by NR treatment, the proportion of rats achieving plantar stepping, parallel paw position, toe clearance and co-ordination were calculated (**Fig. 3E-H**). The improvement in the BBB sub-score was primarily driven by an increase in the proportion of rats that achieved plantar stepping (**Fig. 3E**). Thus, NR treatment successfully increased the hindlimb stepping ability of rats following SCI.

To further assess the extent to which NR treatment improved hindlimb function following SCI, the ability of rats to successfully place their hind paws when crossing a horizontal ladder with irregularly spaced rungs was assessed. The irregular horizontal ladder provides a challenging task for unmasking hindlimb impairments that are not detected when assessing overground locomotion on a smooth surface (Metz and Whishaw, 2002). Successful stepping on the irregular horizontal ladder requires that the rat be able to aim the paw for the appropriate rung, accurately place the paw on the rung and then successfully grasp the rung. This complex task assesses sensory-motor integration and fine motor control and detects deficits in cortical, subcortical and pyramidal systems (Metz and Whishaw, 2002) – axonal systems whose axons may contribute to the spared white matter observed following NR treatment. Beginning two weeks after injury, NR treated rats made fewer missteps in which their hind paw slipped between the rungs (**Fig. 3C**). NR treated rats were also better able to place and reposition their paws on the ladder rungs as indicated by higher ladderwalk scores as determined by the Metz and Whishaw scoring system (8 wpi: NR: 3.9 ± 0.3; control: 2.6 ± 0.3). The horizontal ladderwalk results show that NR treatment improved fine motor control of the hindlimbs following SCI.

## DISCUSSION

There are no clinically approved interventions that protect the spinal cord against injury-induced damage (Resnick, 2013). Moreover, substantial knowledge gaps persist with respect to which biochemical targets can prevent the loss of spinal cord circuitry needed for locomotor function. In other preclinical injury/disease models, one biochemical factor associated with cell death and axon degeneration signaling is NAD^+^ (Alano et al., 2010; Gerdts et al., 2015). In a variety of tissue culture and experimental models, NAD^+^ repletion is neuroprotective and prevents neurodegeneration (Brown et al., 2014; Hou et al., 2018; Vaur et al., 2017) and enhances axon preservation (Araki et al., 2004; Gilley and Coleman, 2010; Sasaki et al., 2006). In experimental SCI, provision of NAD^+^ reduces oxidative stress and neuronal apoptosis (Xie et al., 2017a; Xie et al., 2017b). Collectively, prior studies support the notion that developing therapeutics that maintain or augment NAD^+^ in the early post-injury window may attenuate secondary injury and the propagation of tissue damage.

Until now, the effects of NAD^+^ repletion in established preclinical models of SCI has not been assessed. Several clinical candidates for augmenting NAD^+^ exist, including NR, a naturally occurring vitamin B3. In other preclinical models, NR is protective against neurodegeneration in Alzheimer’s disease (Gong et al., 2013; Hou et al., 2018), excitotoxicity (Brown et al., 2014; Vaur et al., 2017), mitochondrial damage (Khan et al., 2014), peripheral neuropathy (Hamity et al., 2017; Trammell et al., 2016b) and metabolic dysfunction (Canto et al., 2012). NR is available as a nutritional supplement and has been shown to safely elevate NAD^+^ in humans (Airhart et al., 2017; Bogan and Brenner, 2008; Trammell et al., 2016a). These properties of NR have led to interest in using it therapeutically to prevent damage following CNS injury. In the current study, using a preclinical rat thoracic spinal cord contusion model, we tested the effects of NR on SCI repair. The results of this study demonstrate that increasing NAD^+^ levels within the injured spinal cord by administration of NR protects spinal cord tissue from injury and is sufficient to improve motor function following spinal cord injury.

Thoracic SCI in rats and humans results in the formation of a cystic cavity surrounded by a spared rim of white matter (Beattie et al., 1997; Bunge et al., 1993; Hill et al., 2001). In the current study, NR increased the amount of spared spinal cord tissue at the injury epicenter and the penumbra. Analysis of gray matter and white matter sparing revealed that while NR preserved neurons in the gray matter rostral to the injury, its greatest effect was on white matter sparing. This is compatible with the role of NAD^+^ in axon preservation associated with the *Wlds* mutant (Perry et al., 1991; Perry et al., 1990), which has been established to maintain NAD^+^ levels in axons via NMNAT the enzyme responsible for converting NMN to NAD^+^ (Gilley and Coleman, 2010). This also aligns with the neuronal protection observed with NAD^+^ administration following ischemic SCI in rats (Xie et al., 2017a; Xie et al., 2017b) and the protective effects observed in numerous tissue culture and *in vivo* animal studies in which counteracting NAD^+^ depletion protects cells and axons (Liu et al., 2019; Sasaki et al., 2006; Sasaki et al., 2009; Xie et al., 2017a; Xie et al., 2017b).

The extent of functional preservation following SCI at thoracic levels is dependent on the extent of tissue damage (Basso et al., 1996), as well as the location of the white matter damage (Loy et al., 2002a; Loy et al., 2002b). In addition to detecting tissue preservation, in the current study, rats receiving NR treatment showed better locomotor function, as indicated by a reduction of missteps on the horizontal ladder and improvements in hindlimb function as detected by the BBB subscore. This parallels the functional improvements observed 24 h post-ischemic SCI following exogenous administration of NAD^+^ (Xie et al., 2017a; Xie et al., 2017b), as well as the improvements in function achieved with administration of NAM following traumatic brain injury (Goffus et al., 2010) and broadens the conditions in which elevating NAD^+^ is protective for both tissue and nervous system functions.

This study reveals white matter preservation with NR administration after thoracic SCI. In this SCI model, the functional deficits primarily arise from disruption of axonal tracts (Magnuson et al., 1999); future studies defining the specific axonal tracts preserved by NR treatment are needed to understand the underlying circuitry changes that lead to the functional improvements. NR treatment also led to neuron sparing after SCI. The extent to which increasing neurons contributed to motor recovery is less clear. The motor neurons at the thoracic level innervate the T9 intercostal, back and abdominal muscles in a narrow band and the myotome is relatively small. Exploring the functional effects of NR on outcomes after cervical SCI, where the functional deficits are more closely associated with loss of neurons, would be helpful for determining the impact of preserving neurons using NR. It is possible that greater functional benefit following NR treatment could be achieved in an injury where lower motor neuron injury directly impacts the functional deficits.

Due to its critical functions in cells, the cellular biosynthesis of NAD^+^ is highly controlled (Covarrubias et al., 2021). The enzymes responsible for generating NAD^+^ have tissue specific distribution (Mori et al., 2014). Examination of available gene expression databases for the rodent spinal cord showed that the key enzymes needed to synthesize NAD^+^ from NR are expressed within the spinal cord. The expression of *Nmrk1* and *Nmnat1–3*, along with the quantification of spinal cord NAD^+^ showing its elevation in the spinal cord in response to NR administration supports the premise that following NR administration NAD^+^ is synthesized in spinal cord cells. Confirmation of protein expression of the enzymes responsible for synthesizing NAD^+^ within the spinal cord following injury would lend further support to this premise.

A variety of NR doses and delivery methods have been used to test its effects in rodents (Brown et al., 2014; Canto et al., 2012; Gong et al., 2013; Hou et al., 2018; Khan et al., 2014; Trammell et al., 2016a; Trammell et al., 2016b; Vaur et al., 2017; Zhang et al., 2016). Doses have ranged from 200 mg/kg in experiments assessing its functional effects on peripheral neuropathy using oral gavage (Hamity et al., 2017) to 1000 mg/kg i.p. twice a day in experiments testing its effects in noise-induced hearing loss (Brown et al., 2014). The dose of NR used in the current study is similar to those in several previous studies (Canto et al., 2012; Khan et al., 2014; Trammell et al., 2016b; Zhang et al., 2016). Furthermore, administration of 500 mg/kg i.p. to rats raised peripheral NAD^+^ levels in the liver and skeletal muscle to levels reported previously in mice following oral administration of NR via chow (Canto et al., 2012).

Compared to peripheral tissues (Gong et al., 2013; Hou et al., 2018; Vaur et al., 2017), CNS elevation of NAD^+^ following NR administration is substantially lower. In the injured spinal cord, the increase in NAD^+^ achieved with 500 mg/kg of NR, albeit protective, was relatively modest (2-fold). In models of brain injury and disease, a similarly modest increase in NAD^+^ also correlates with tissue protection with NAD^+^ manipulating therapies (Gong et al., 2013; Hou et al., 2018; Vaur et al., 2017; Zhao et al., 2021). This implies that a large increases in NAD^+^ levels may not be needed to support NAD^+^-dependent cellular activities in the CNS. There does appear to be a minimal level of NAD^+^ augmentation required for spinal cord protection. Experiments testing a lower dose of NR (250 mg/kg) it failed to elicit SCI repair (unpublished data). Further studies are needed to define the NR dosing conditions and the level of NAD^+^ augmentation necessary for maximal SCI repair with NR-based therapies. Additionally, NR was administered starting 4 days prior to SCI, following the rationale that *in vitro* preincubation with compounds that alter NAD+ levels/synthesis is needed for axon protection in some models (Sasaki et al., 2006). Thus, it will be important to establish whether similar protection is achievable if NR is administered only post-SCI.

After SCI, elevated levels of NAD^+^ were not detected in the brain following NR administration. This aligns with prior studies examining metabolic and muscle defects and NR bioavailability where increased NAD^+^ levels were not detected in the brain (Canto et al., 2012; Khan et al., 2014; Trammell et al., 2016b). This could reflect impaired entry for NR into the CNS. In a preliminary dose-response study using naive rats, we were able to detect elevated levels of NAD^+^ in the brain. Thus, it seems more plausible that the difference in NAD^+^ levels detected in the brain between naive and SCI-rats reflects a difference in NAD^+^ consumption between healthy vs stressed/damaged tissue. The enhancement in NAD^+^ in the spinal cord, but not the brain, following SCI could reflect increased entry of NR into the spinal cord due to disruption of the blood-spinal cord barrier (BSCB) during the administration window post-SCI (Matsushita et al., 2015). Studies examining NAD^+^ consumption and/or visualizing the biodistribution of NR within the CNS post-SCI would be helpful in determining the cause of the difference in the NAD^+^ levels achieved within the brain and spinal cord.

NR offers several advantages over other NAD^+^ precursors that may make it more suitable as a therapeutic (Elhassan et al., 2017; Trammell et al., 2016a). Lower concentrations of NR are required compared to NMN to obtain a similar increase in NAD^+^ (Trammell et al., 2016a) and it does not cause niacin toxicity, like skin flushing, as NMN does (Elhassan et al., 2017). However, a substantial proportion of NR is converted to nicotinamide (NAM) upon entry into the bloodstream following oral or intraperitoneal administration of NR (Cercillieux et al., 2022; Liu et al., 2018), which could contribute to the relatively low levels of NAD^+^ achieved within the CNS. Newer NR derivatives with chemical modifications are in the pipeline (Makarov and Migaud, 2019; Zarei et al., 2021). Dihydronicotinamide riboside (NRH), a newly identified NAD^+^ precursor, appears to be more potent at increasing NAD^+^ than other precursors (Giroud-Gerbetant et al., 2019; Yang et al., 2019). therefore, these novel NR derivatives could overcome some of the limitations of raising NAD^+^ levels, particularly within the CNS.

The mechanism(s) by which increasing NAD^+^ with NR leads to spinal cord repair is not examined in this study. Further experiments are required to determine how NR increases NAD^+^ in the spinal cord and leads to tissue protection. By supplementing NAD^+^, NR may circumvent the post-injury depletion of NAD^+^ caused by enhanced activity of the NAD^+^-consuming enzymes (e.g., PARPs, CD38/157, Sirtuins, SARM1) that are important for DNA repair, mitochondrial biogenesis and function, calcium signaling, and axon injury induced degeneration (Covarrubias et al., 2021; Elhassan et al., 2017; Gerdts et al., 2016). Additionally, they have been linked to cellular changes associated with spinal cord tissue damage, including neuronal cell death, neuroinflammation, glial activation, and neurodegeneration and impaired axon regeneration in conditions of demyelination (Liu et al., 2021; Roboon et al., 2019; Scott et al., 1999). Thus, increasing NAD^+^ levels through NR administration could facilitate the maintenance of cellular energy production (Khan et al., 2014; Trammell et al., 2016a; Trammell et al., 2016b), thereby preventing cell death and secondary injury associated with ATP depletion present after SCI (Anderson et al., 1982; Braughler and Hall, 1984; Wu et al., 2009).

Application of NAD^+^ precursors can suppress PARP and CD38 associated cellular changes (Roboon et al., 2019; Scott et al., 2004). Reducing consumption of NAD^+^ by PARP-1 and CD38 increases NAD^+^ bioavailability for the sirtuins, which have a higher Km for NAD^+^ (Bai et al., 2011; Barbosa et al., 2007) and are associated with NR-mediated tissue protection (e.g., (Brown et al., 2014)). The sirtuins deacetylate (a post-translational protein modification) specific cellular proteins that modulate a variety of metabolic, energetic and stress, and transcriptional response pathways in cells, and are linked to tissue protection after SCI (Chen et al., 2017; Lin et al., 2021). Characterization of the activity of these NAD^+^ consuming enzymes within the spinal cord following SCI could not only be insightful in terms of understanding their contribution to NR-mediated protection, but could lead to the identification of their role(s) following SCI and the development additional ancillary therapeutics to specifically modify their activity.

## CONCLUSIONS

NR is a safe and effective strategy to enhance NAD^+^ levels (Airhart et al., 2017; Bogan and Brenner, 2008; Covarrubias et al., 2021; Trammell et al., 2016a). The current study demonstrates that NR treatment enhances tissue preservation and promotes recovery of function following SCI. It supports the need for further testing of NR as a therapy for SCI. Additional mechanistic studies are needed to establish the specific mechanism by which the elevation of NAD^+^ by NR administration leads to protection following SCI. Examination of the contribution of the various NAD^+^ consuming enzymes during the acute and subacute injury intervals could lead to a better understanding of how NAD^+^ is depleted following SCI and how these enzymes contribute to tissue destruction following SCI.

## ACKNOWLEDGEMENTS

Dr. Anthony Sauve provided the nicotinamide riboside and contributed to the initial conceptualization of the project. The authors would like to thank: Jaime Tanner, Elena Chepurko and Jennifer Brown for their assistance with animal care, behavior and tissue collection; and Sarah Groover and Abhay Deshmukh for assistance with tissue processing and quantification.

